# EpCAM Aptamer siRNA chimeras: Therapeutic efficacy in epithelial cancer cells

**DOI:** 10.1101/656199

**Authors:** Jayashree Balasubramanyam, Lakshmi Badrinarayanan, Bharti Dhaka, Harsha Gowda, Akhilesh Pandey, Krishnakumar Subramanian, Lakshmi B. Subadhra, Sailaja V. Elchuri

## Abstract

In the era of personalized medicine as well as precision medicine, targeted therapy has become an integral part of cancer treatment in conjunction with conventional chemo- and radiotherapy. We designed aptamer-siRNA chimeras that can specifically target cancers expressing EpCAM, a stem cell marker and deliver the specific siRNA required for therapy response. The siRNAs were chosen against PLK1, BCL2 and STAT3 as these oncogenes play prominent role in tumour progression of several cancers. Targeted delivery of EpCAM-siRNA chimeras resulted in cell death in several cancer cell lines such as cancers of the breast, lung, head and neck, liver and retinoblastoma. *In vivo* analysis of EpCAM-siRNA chimera mediated silencing on RB xenografts tumour model showed increased tumor reduction in all the three EpCAM-siRNA treated conditions. However, regulation of PLK1 exhibited higher efficacy in tumour reduction. Therefore. We studied signaling mechanism using global phosphoproteomics analysis. An increased P53 mediated downstream signalling pathway might have enabled increased apoptosis in the cancer cells. In conclusion, this study demonstrated the efficacy of EpCAM aptamer chimeras coupled to siRNA gene silencing for targeted anti-cancer therapy.

**Graphical abstract:** 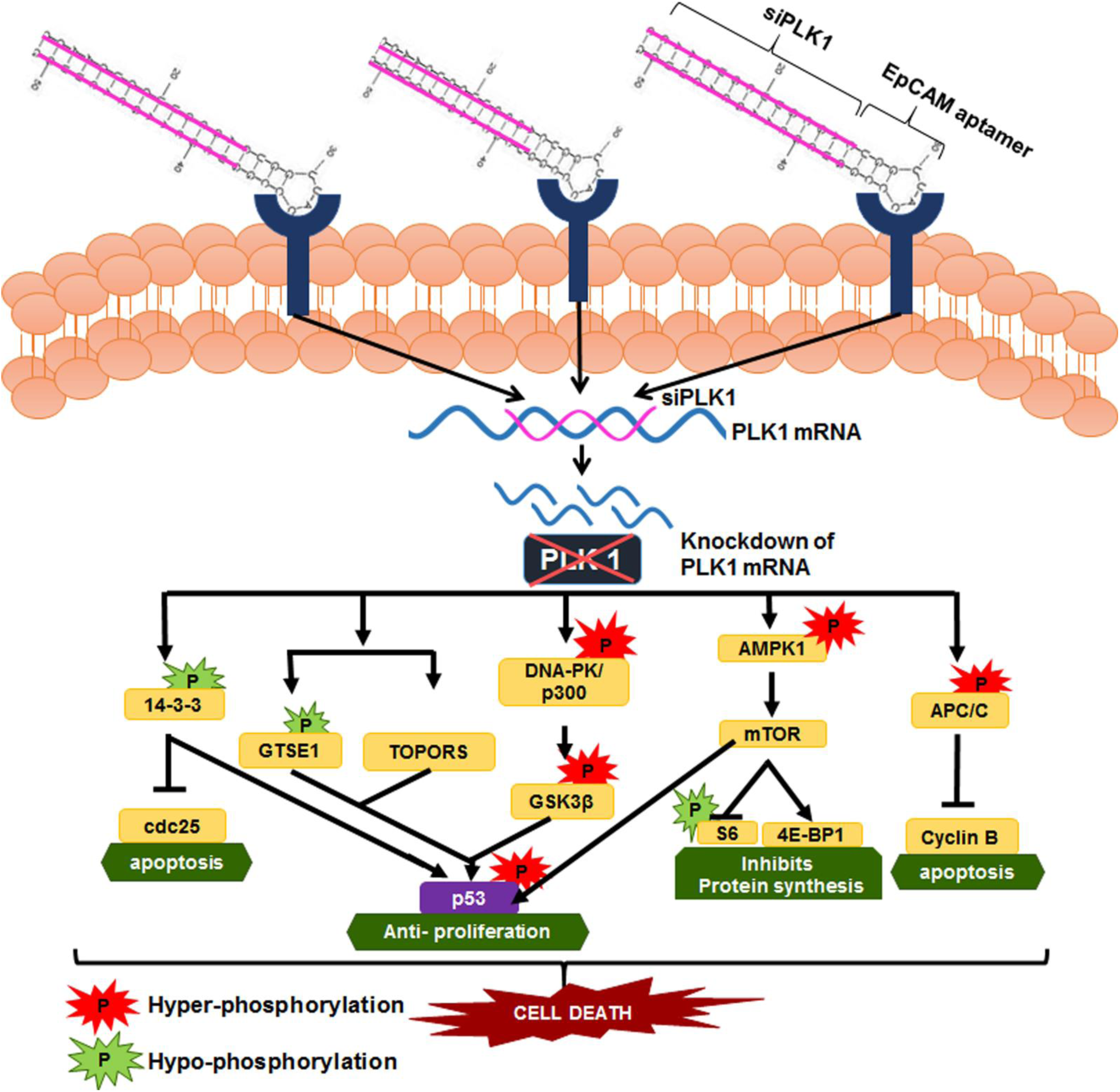

Illustration showing how EpCAM aptamer-mediated silencing of PLK1 could control the cell cycle progression at multiple number of check points and induce apoptosis involving hyper and hypophosphorylation of variety of signalling molecules

## INTRODUCTION

Limitations of conventional chemotherapeutic drugs due to non-specificity present a major challenge for cancer therapy. These shortcomings are largely attributed to the ability of cancer cells to repopulate and metastasize after initial therapies which makes them resistant to chemotherapy and radiotherapy. Thus there is a need to eliminate the cancer cells by specifically targeting the mechanisms which promote the cells survival and progression. The most common method to hamper the cell cycle progression is by blocking the activity of selected genes using a siRNA. Short interfering RNAs (siRNAs) are oligonucleotide-based pharmaceuticals with a promising role in targeted cancer therapies ^1^. These have great therapeutic potential and specificity, but their shortcomings remains the delivery and internalisation into cells and off-target effects. Though lipid based delivery of siRNA is routinely used, its efficiency is hampered due to its *in vivo* instability and lack of target tissue delivery. Numerous strategies are underway to eliminate cancer cells specifically by targeting surface biomarkers coupled with regulation of oncogenes and application of novel delivery technology to combat cancer. Hence, drug delivery systems using aptamer technology are used to enhance the specific delivery of these siRNAs to the targeted sites ^2^. Aptamers are short RNA/DNA molecules that specifically bind to target proteins with high affinity ^3, 4^. Various forms of aptamer-chimeras, (aptamers conjugated with drugs and biomolecules like siRNAs) have been extensively applied in many cancer therapeutics ^5^. These aptamers not only bind to specific cancer cells but are also internalized by the process of receptor mediated endocytosis and deliver the respective biomolecule attached to it^6^. Aptamer-siRNA chimeras have been found to be proficient in inhibiting tumour progression^7^. Various targeting aptamers have been reported till now, which include prostrate specific membrane antigen (PSMA)^7^, B cell activating factor receptor (BAFF-R)^8^ and Epithelial Cell Adhesion Molecule (EpCAM)^6^. These targets are mainly chosen depending on their exclusive expression on the specific cancer cell type. EpCAM is a well-established cancer stem cell marker identified in many tumours. A nuclease-resistant aptamer targeting EpCAM was found to be clinically beneficial for both diagnostics and therapeutics ^9^. EpCAM aptamer chimera conjugated with PLK1 siRNA was also found effective in restricting tumour growth in breast cancer ^10^. However studies are lacking on the efficacy of these EpCAM aptamer chimeras in reducing tumour growth in other cancer models. Hence, targeting various cancer cells using EpCAM aptamer-siRNA chimera against oncogenes that are overexpressed in several solid tumours would be an effective anticancer therapeutic in combating different cancers.

Many anticancer drugs have been developed targeting oncogenes due to their ability to suppress cell death such as Polo-like Kinase 1 (PLK1), B-cell Lymphoma 2 (BCL2) and Signal transducer and activator of transcription 3 (STAT3)^11, 12^. PLK 1 is a critical oncogene that is involved in centrosome maturation in the late G2/early stages of cell cycle progression^13^ and activation of Cdc25 ^14^, thereby leading to uncontrolled cell division and cancer progression. While PLK1 contributes to cancer by inducing uncontrolled cell progression, BCL2 acts as anti-apoptotic marker by preventing cell death^15^. Overexpression of BCL2 has been seen in many cancers especially in invasive tumours and is known to play a major role in the survival of the cell ^16, 17^. An aberrant STAT3 signalling promotes initiation and progression of many cancers either by inhibiting apoptosis or by inducing cell proliferation ^18^. Furthermore, consistent overexpression of STAT3 was observed in angiogenesis, cellular invasion and survival ^19^. Thus targeting these oncogenes and blocking their expression would inhibit cell proliferation and result in reduced tumour growth. In our previous study we have shown that EpCAM aptamer-siPLK1 chimera identified a novel prominent role of lipid synthesis for the oncogene PLK1 ^20^. However, extensive investigations on reducing tumour burden and signalling mechanism leading to cancer cell death was not studied. Additionally, efficacy of EpCAM aptamer mediated regulation of BCL2 and STAT3 are not reported before. Hence, we designed and studied the EpCAM aptamer chimeras with siRNA for PLK1 (EpCAM-siPLK1), BCL2 (EpCAM-siBCL2) and STAT3 (EpCAM-siSTAT3) in several cancers which overexpress EpCAM. The *in vitro* cell line model that exhibited superior efficacy was further tested in mouse xenograft model. The molecular signalling mechanism was studied for the best performing aptamer-siRNA chimera using SILAC-based phosphoproteomics analysis.

## RESULTS

We targeted various cancer cells specifically by using an already available EpCAM binding RNA aptamer. This served as a carrier molecule to deliver the siRNAs (siPLK1, siBCL2, siSTAT3) to EpCAM expressing cancer cells.

### Designing of EpCAM-siRNA chimeras

EpCAM aptamer chimeras were designed using bioinformatics approaches. The silencing activity and specificity of these aptamer-siRNA chimeras were enhanced by incorporating few modifications like adding 2-nucleotide, 3’-overhangs to increase its stability against endogenous cell nucleases and including a dicer recognition site. The aptamers were designed having the sense strand of the siRNA starting at the 5’ end of 20 nt length followed by the truncated EpCAM aptamer sequence of 20 nt and then the antisense strand having 20 nt with a 2 nt (UU) overhang a the 3’ end. This design increases the stability of the chimera and makes it easy for the dicer enzyme to cleave the chimera at the 21^st^ nucleotide and release the siRNA from the EpCAM aptamer (**Figure 1A**). The EpCAM aptamer sequence is in the middle of the sense and antisense strand in each EpCAM-siRNA chimera. The 3’ end of the chimera is fluorescently labelled with FITC. The designed aptamer was cleavable by dicer enzyme *in vitro* into EpCAM aptamer and the respective siRNA (**Figure 1B**).

**Figure 1:**
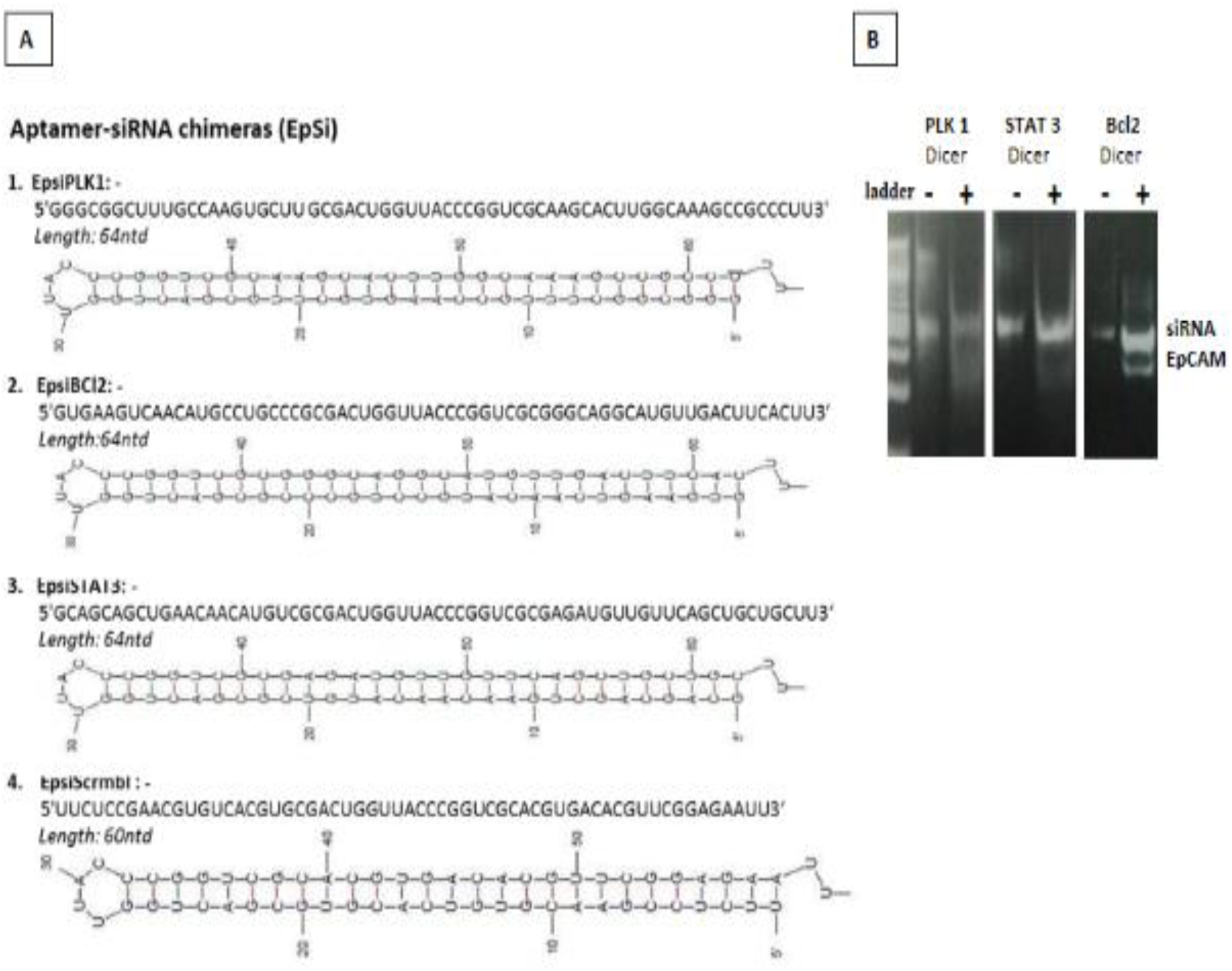
(A) Predicted secondary structures for the EpCAM-siRNA chimeras. (a) EpCAM-siPLK1, (b) EpCAM-siSTAT3, (c) EpCAM-siBCL2 and (d) EpCAM-siScr. The aptamer region is truncated and is represented with a black outline continued by the 20-nt siRNA duplex. **(B) Dicer assay**. Chimeras were treated with human recombinant dicer for 6 h. the digested samples were resolved with 3% agarose gel electrophoresis. Two bands, EpCAM and siRNA, were produced in the presence of dicer enzyme (dicer +).

### Expression of EpCAM in different cancer cell lines

Several cancer cells such as breast, pancreatic, lung cancer, head and neck and Retinoblastoma (RB) were examined for the expression of EpCAM using PCR and quantified using qPCR (**Figure 2**). The presence of EpCAM was found to differ in each cancer cell line in comparison with a well-known EpCAM negative (EpCAM^−^) Muller glial (Mio M1) normal cell line. Mio M1, WERI Rb 1, Y79 (RB cell lines), MiaPaCa 1, PanC1 (Pancreatic cancer cell lines), A549, Ncl H23 (Lung cancer cell lines) and HUH-7 (Hepatocellular carcinoma cell line) showed very minimal or even negligible expression of EpCAM (**Figure 2, lane 1,3,4,6,7,8,9,11**). Whereas, MCF-7 (Breast cancer cell line), NCC RbC 51 (RB), HepG2 (Hepatocellular carcinoma) and RPMI 2650 (Head & neck) cancer cell lines showed high EpCAM expression levels (**Figure 2, lane 2,5,10,12**). Based on these results, the cell lines were categorised into EpCAM^**+**^ and EpCAM^−^ cell lines. The efficacy of aptamers was tested in EpCAM^**+**^ cell lines compared to Mio M1 as EpCAM^−^ control cell line.

**Figure 2:**
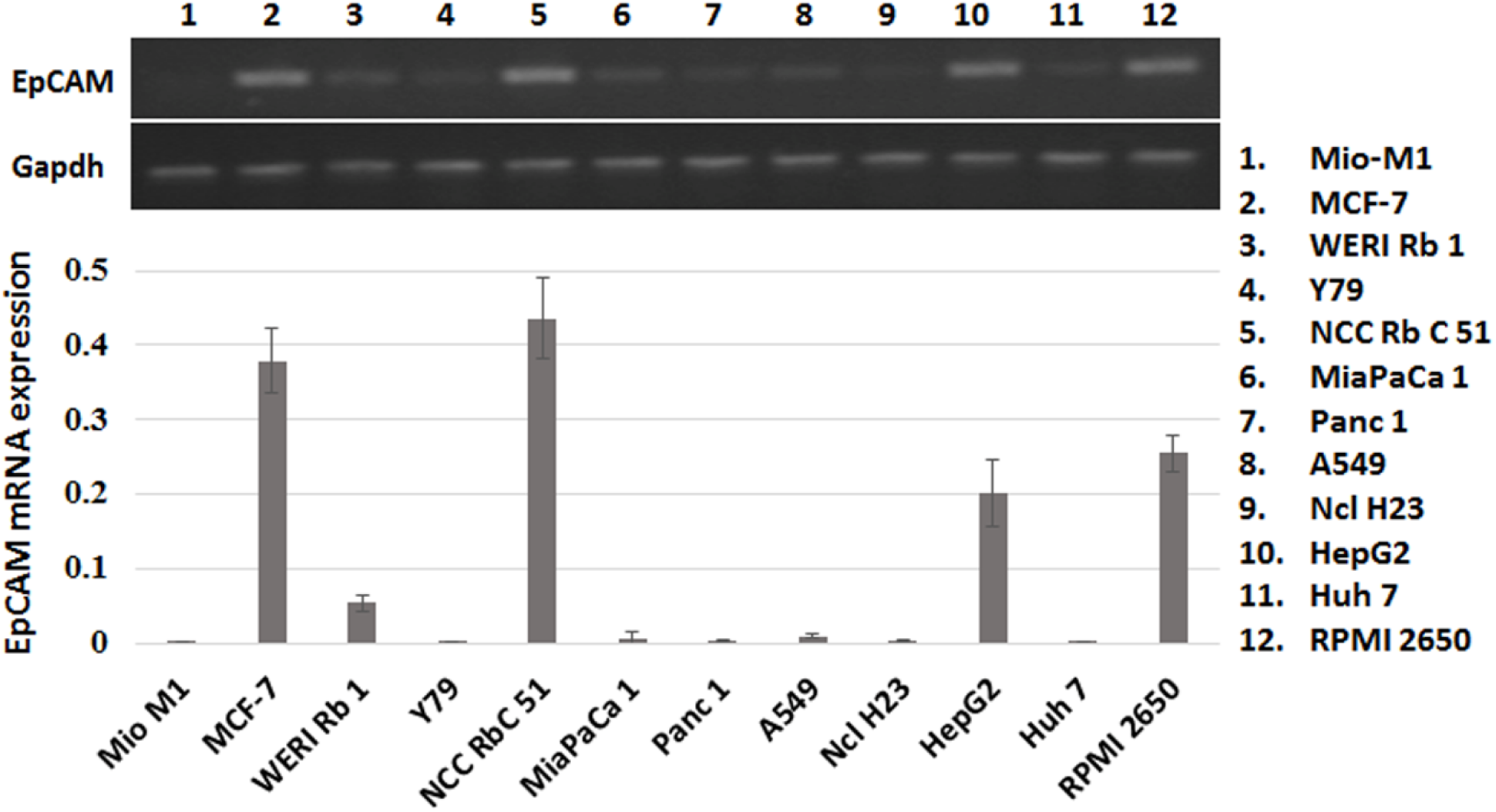
Native expression of EpCAM gene in different epithelial cancers. The native expression of EpCAM was assessed in 12 different epithelial cancer cell lines using PCR and resolved on 3% agarose gel electrophoresis. The lanes represent (1) Mio M1, (2) MCF-7, (3) WERI Rb 1, (4) Y79, (5) NCC RbC 51,(6) MiaPaCa 1, (7) Panc 1, (8) A549, (9) Ncl H23, (10) HepG2, (11) Huh 7, (12) RPMI2650. Significant increased levels of EpCAM expression were seen in MCF-7, NCC RbC 51, HepG2 and RPMI 2650. Data are means ± SEM, *n*=3.

### Binding and internalisation of the chimeras in EpCAM expressing cell lines

The binding and internalisation of the EpCAM chimeras in various epithelial cell lines were compared in those cell lines which showed higher amount of EpCAM expression. These chimeras were designed specifically to bind to EpCAM^+^ cells and deliver the siRNA. Since the aptamers are FITC labelled, its binding efficacy in EpCAM expressing cell lines was examined by flow cytometry (**Figure 3A**) and fluorescence microscopy (**Figure S1**). As shown in **Figure 3A**, the fluorescence intensity from the chimeras bound to NCC RbC 51 and MCF-7 was higher than those bound to HepG2 and RPMI 2650. However, all cell lines exhibited greater binding efficiency compared to the control cell line Mio M1 which showed very minimal binding (10%) even at a very high concentration (1000nM) of the chimera. About 90% of binding was achieved in NCC RbC 51 with a minimum concentration of 100nM EpCAM-siRNA. In MCF-7 cells, as the concentration of the chimera increased, a gradual increase in binding was seen. 85% binding was seen with 300 nM aptamer concentration. However, further increase in the concentration resulted in saturation of binding to MCF7 cells. Whereas, in RPMI 2650 and HepG2 cells only 50-60% binding of EpCAM-siRNA was observed even at a very high concentration of 500nM. These results indicate that the binding of the chimeras to the cancer cells may be proportional to the magnitude of EpCAM expression in these cells. Furthermore, higher expression of EpCAM on MCF-7 and NCC-RBC 51 might have enabled higher aptamer chimera binding on these cells.

**Figure 3:**
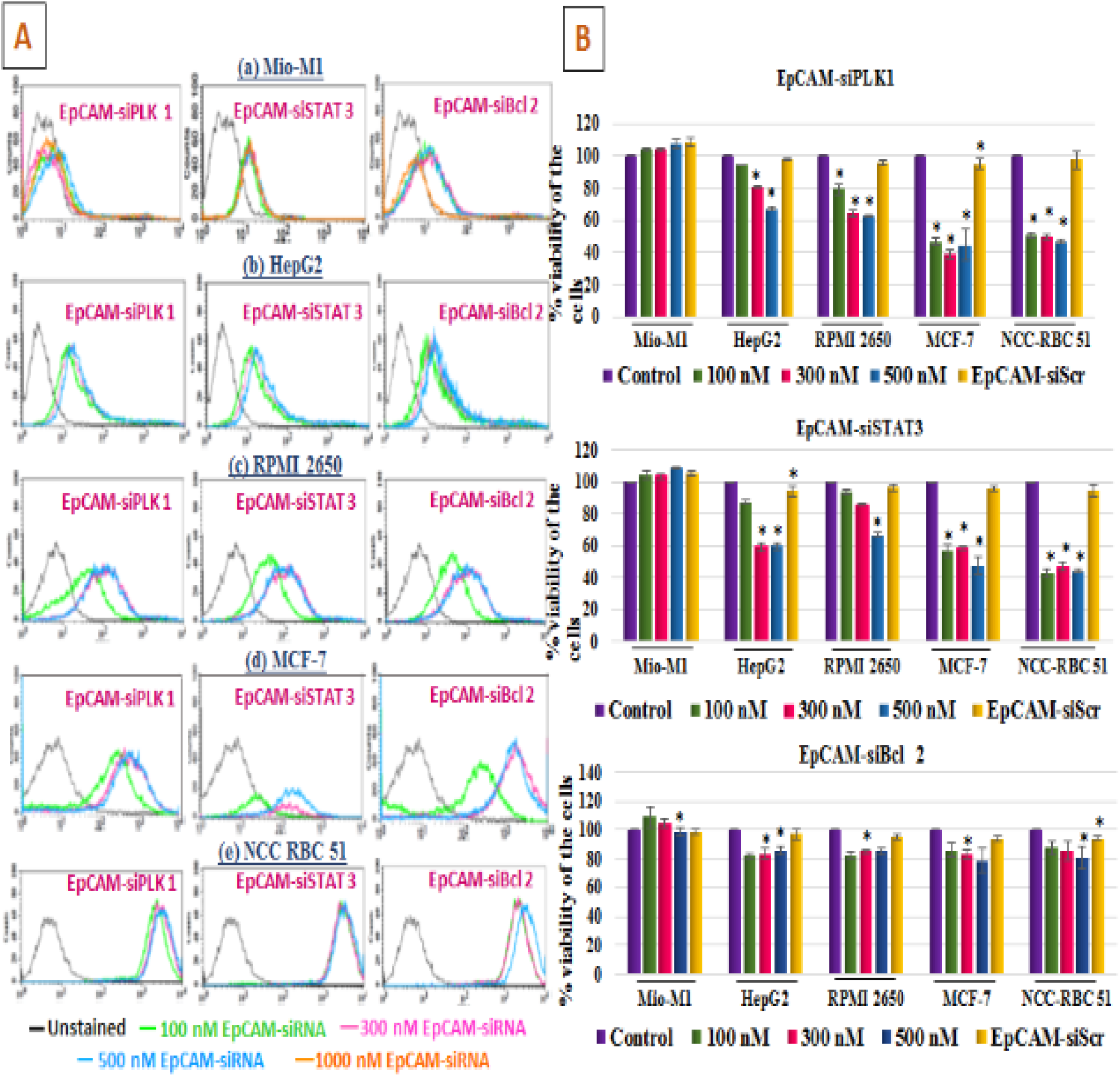
Aptamer-siRNA chimeras bind specifically to the cell surface antigen EpCAM and cause cell death. (A) Binding profiles using Flow Cytometry (a) Mio M1 (b) HepG2 (c) RPMI 2650 (d) MCF-7 and (e) NCC RbC 51 were incubated with FITC labelled EpCAM-siRNA chimera. Cell surface binding of the chimeras at different concentrations (100 nM in green, 300 nM in pink, 500 nM in blue and 1000 nM in orange) are represented as flow cytometric overlay plots. (B) Cells were assessed for percentage viability after different concentrations of EpCAM-siRNA treatment (100 nM in green, 300 nM in pink, 500 nM in blue and EpCAM-siScr in yellow) using MTT reagent. The reduction in the number of viable cells showed that cell death has increased after the treatment with EpCAM-siRNA chimeras. Maximum binding efficiency and cell death was seen in NCC RbC51 and MCF-7 cell lines with 100 nM of treatment. Data are means ± SEM, *n*=3. *,*P*< 0.05; (as denoted).

### EpCAM-siRNA mediates dose dependent cytotoxicity

To assess the cytotoxic effect of EpCAM-siPLK1, EpCAM-siSTAT3 and EpCAM-siBCL2, a panel of cancer cell lines and control cell line were exposed to the chimeras for 72 h at varying concentrations ranging from 100-500 nM and cell viability was assessed using MTT assay (**Figure 3B**). A very minimal cytotoxicity was seen in the control cell line Mio M1 after treatment with all the three chimeras. These aptamer chimeras could not bind to the Mio M1 cells **(Figure 3B)**, resulting in lack of cell death in these cells. In RPMI 2650 and HepG2 significant (p value < 0.05) cell death was observed only at higher concentrations of Ep-siPLK1 and Ep-siSTAT3 (500nM). This is in agreement with the EpCAM expression in both RPMI 2650 and HepG2 where moderate expression of EpCAM was observed **(Figure 3B)**. The expression of moderate levels of EpCAM moieties could have necessitated greater amounts of the aptamer chimeras needed for cancer cell death. However, in MCF-7 and NCC RbC 51, cells a significant cytotoxicity (p< 0.05) was observed at 100 nM concentration due to the abundant expression of EpCAM in these cells. The magnitude of decrease in cell death was lower in all the cell lines on treatment with EpCAM-siBCL2 compared to EpCAM-siPLK1/ EpCAM-siSTAT3 aptamer chimeras. Despite showing high percentage of binding in FACS assay, it showed only a moderate amount of cell death in all the cell lines. Cytotoxicity was seen in EpCAM^**+**^ compared to EpCAM^−^ MIO M1cells, indicating the specificity of EpCAM expression mediated targeted cell death. Additionally, cell death was minimal in EpCAM-Scr aptamer treated cancer cells indicating non-toxicity of the EpCAM aptamer.

### EpCAM aptamer mediated cell type specific knockdown of target genes

Next we studied the ability of the aptamer chimeras to silence the expression of the chosen oncogenes specifically in EpCAM^**+**^ cancer cells. The chimera once specifically bound to EpCAM positive cells, can internalise and deliver the respective siRNA tagged to it, which binds to the target gene and silence its expression. The three siRNAs used in our study are specific to important oncogenes which, when silenced can result in inducing apoptosis and eventually causing cell death. The minimal concentration of the chimeras which showed maximum cytotoxic effect for the specific cell line were chosen for further analysis A concentration of100 nM was used for MCF-7 &NCC RbC 51 wheras 500 nM was used for HepG2 and RPMI 2650. The reduction in PLK1, STAT3 and BCL2 gene expression on treatment with the EpCAM-siRNA chimeras were compared to siRNA alone to investigate the targeted ability of these aptamers in EpCAM^+^ cancer cells. In the EpCAM^**−**^ cell line Mio M1, the knockdown of PLK1, BCL2 and STAT3 were observed in siRNA treated cells alone and not in the aptamer chimera treated cells. In RPMI 2650 and HepG2 cells, the downregulation of the oncogenes was higher in siRNA treated cells compared to their aptamer chimera treated cells (**Figure 4A**). The partial knockdown in the oncogenes observed in HepG2 and RPMI 2650 could be due to moderate binding of the aptamer chimeras regulated through the EpCAM expression in these cells. Whereas in MCF-7 and NCC RbC 51 the downregulation of PLK1, BCL2 and STAT3 genes were significantly (P> 0.05) not different in both EpCAM-siRNA and siRNA treated cells indicating that lipofectamine delivers the siRNA to all the cells, whereas, the aptamer chimera delivered the siRNAs to EpCAM^+^ cells (**Figure 4A)**. The superior functional performance of the aptamer chimeras in NCC RbC 51 cell line enabled us to choose this *in vitro* tumour cell model for further experiments.

**Figure 4:**
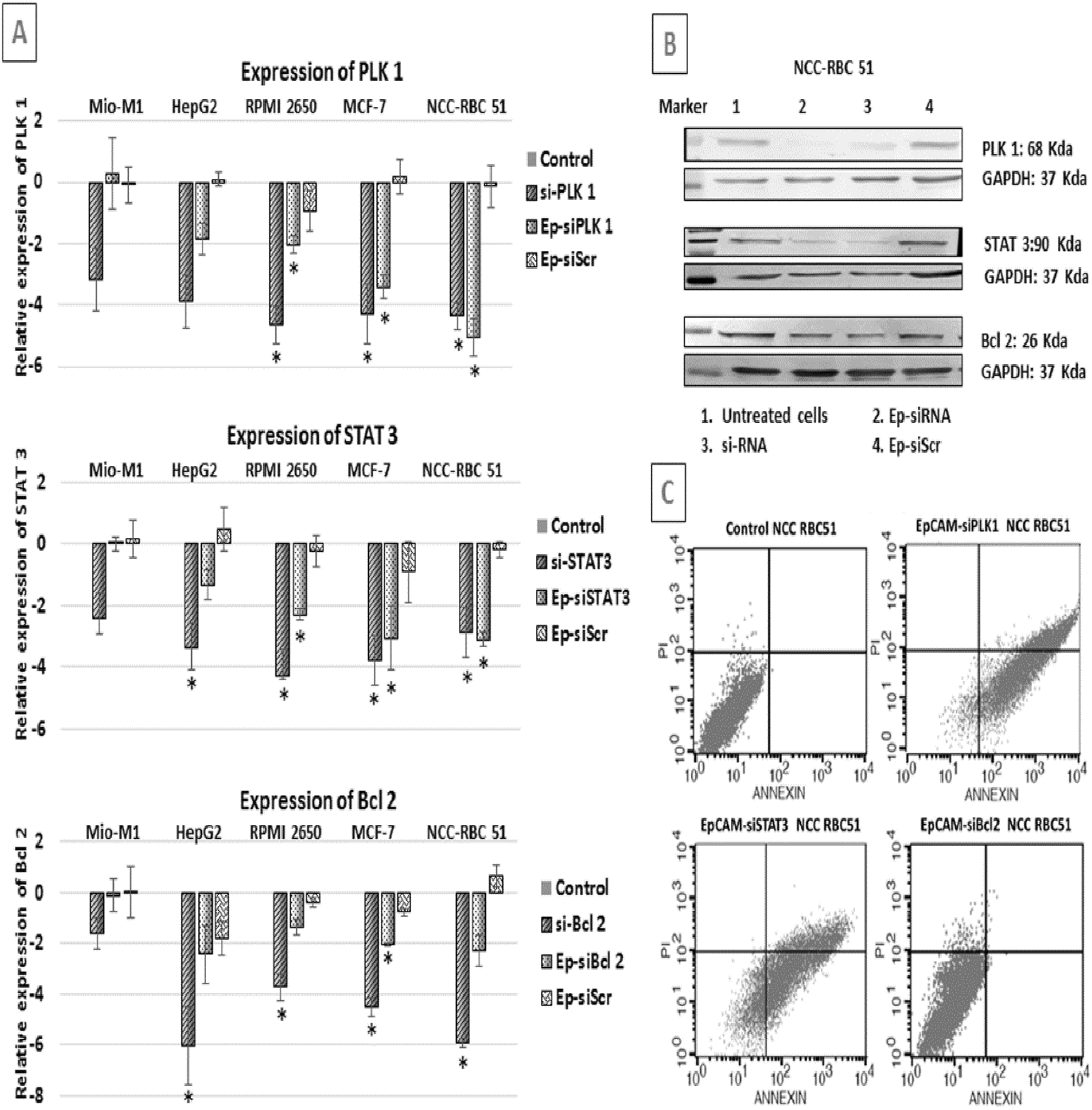
Cell-type specific silencing of genes with aptamer chimeras and their apoptotic effect. Cells incubated with various aptamer chimeras for 24h in the absence of transfection reagents (a) silencing of PLK1 (b) BCL2 (c) STAT3 expression was assessed by (A) quantitative PCR for all the cell lines and (B) western blot for NCC RbC51. Efficient Gene silencing due to EpCAM-siRNA was restricted to NCC RbC51 cells and correlated with efficient binding of the chimera in these cells. (C) Flow cytometry showed the increased populations at early (Annexin V+PI^−^) and late (Annexin V+PI^+^) apoptosis. The stage movement shows that apoptosis occurs upon EpCAM-siRNA treatment. Data are means ± SEM, *n*=3. *,*P*< 0.05; (as denoted).

The knockdown of PLK1, STAT3 and BCL2 at protein level was checked in NCC RbC 51 cells after treatment with the aptamer chimera. The downregulation of PLK1 and STAT3 were found to be more in EpCAM-siRNA chimera treated cells compared to siRNA treated cells (**Figure 4B**). The siRNA was more effectively delivered by EpCAM aptamer than lipofectamine, which shows that this could be a potential targeted delivery system for EpCAM positive cancers. However, magnitude of BCL2 protein reduction was lowest even though the binding of the chimera to the cancer cells was same as other two chimeras. The EpCAM-siScr chimera did not reduce oncoprotein expression. Therefore, these results show that the EpCAM-siRNA, but not EpCAM-siScr, is able to deliver the siRNA and regulate the gene/protein expression in a cell type specific manner.

### EpCAM-siRNA induced apoptosis in NCC RbC 51 cell line

To confirm the occurrence of apoptosis on treatment with EpCAM-siPLK1, EpCAM-siSTAT3 and EpCAM-siBCL2 in NCC RbC 51, Annexin-PI staining was performed and analysed using flow cytometry after 48 h treatment with EpCAM-siPLK1 (**Figure 4C**). There was an increase in early apoptotic cell population (Annexin V+ PI^−^) from 0.34% to 39.00% and an increase in late apoptosis (Annexin V+ PI^+^) from 0.12% to 47.02%. We did not observe any necrotic population. EpCAM-siSTAT3 treatment, resulted in an increase in the early apoptotic cell population (Annexin V+ PI^−^) from 0.47% to 15.99% and an increase in late apoptosis (Annexin V+ PI^+^) from 0.22% to 22.35%. Additionally, the cells in the process of necrosis increased from 1.12% to 19.6%. However, consistent with lower functional activity of EpCAM-siBCL2, reduced cell death through apoptosis was observed in NCC RbC 51 cells (Figure 4C). About 88.38% of cells were found to be live, whereas 1.41% in early apoptotic, 2.49% in late apoptotic stage. However, 5.91% cells were observed in necrotic stage. The shift of the cells towards apoptotic stage indicates that the treatment with EpCAM-siRNA aptamers supresses the proliferation of NCC RbC 51 cells after the regulation of oncogenes by aptamer chimeras in this cell line.

### Reduction of tumour burden through systemic administration of EpCAM-siRNA chimeras in the mouse xenografts model of RB

To assess the effect of the EpCAM-siRNA chimeras on tumour growth *invivo*, NCC RbC 51cells were injected subcutaneously in the right flank of the animal and tumour xenografts were established in athymic Nude-Foxn1nu male mice. Once the tumour attained a palpable size, animals were randomized based on tumour volume (TV ≈ 200 mm^3^) and dosing was initiated. EpCAM-siRNA (100 μl, 0.8 nmol per mouse) was injected intraperitoneally to tumour bearing mice every third day for 30 days (10 cycles).

A significant tumour reduction (p<0.05) was seen in EpCAM-siPLK1 and EpCAM-siSTAT3 treated mice when compared to the control (**Figures 5A and B**). Howerver, a greater percentage of tumour reduction (56.11%) was seen in EpCAM-siPLK1 treated mice compared to aptamer chimeras after 10 cycles of treatment in a span of 30 days. The tumour size was nearly 1600mm^3^ in control and reduced to 834 mm^3^ in EpCAM-siPLK1 and 988 mm^3^in EpCAM-siSTAT3 chimeras treated mice, respectively. However, when treated with EpCAM-siBCL2, the average tumour size was found to be 1250 mm^3^ showing a reduction of only 34.21% compared to vehicular control mice. Upon treatment with EpCAM-siBCL2, a significant (p<0.05) reduction was observed in 3 animals, whereas, 2 animals were found to be non-responders indicating reduced ability of this chimera to arrest tumour growth compared to other two chimeras. However, all the three aptamer chimeras were non-toxic to the animals. Furthermore, the body weight of the animals was also found to increase in a definite manner in both treated and untreated conditions till the end of the study (**Figure 5C**). Additionally, the aptamer chimera treatments did not cause any structural changes in the detoxifying organs such as liver and kidney **(Figure S2)**.

**Figure 5:**
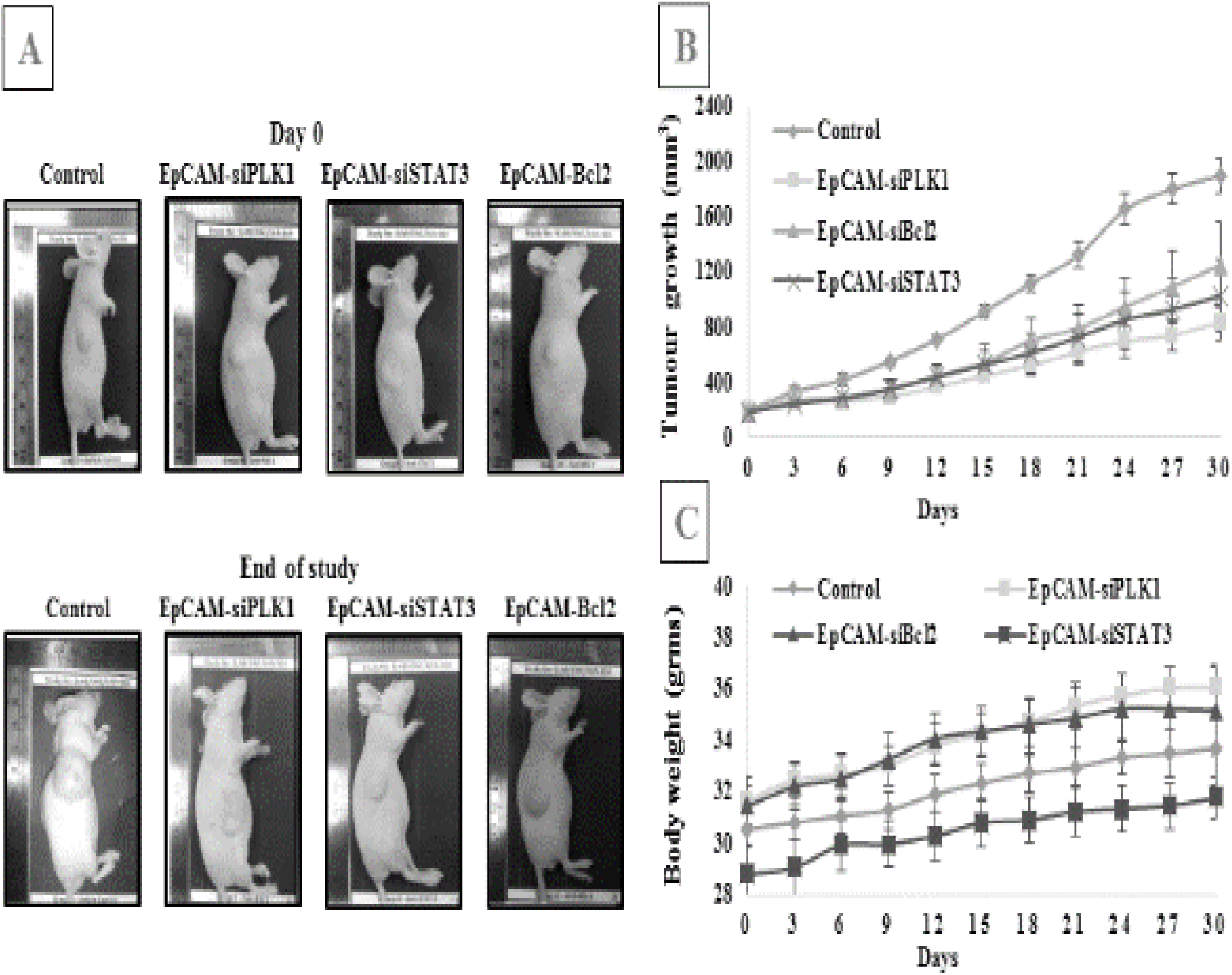
Administration of EpCAM-siRNA reduces tumour growth in a xenograft model of RB. Chimeric RNAs were administered intraperitoneally in mouse xenograft model bearing NCC RbC 51 RB cancer cells implanted into the subcutaneous flanks of the nude mice. Saline treated animals were used as control. (A) Representative tumour-bearing mouse images: control and EpCAM-siRNA treated conditions. (B) Corresponding tumour growth curves. After treatment, tumours were measured every third day (n=5, p<0.05). (C) Body weights were measured after the treatment. A gradual minimal increase in body weight was seen in the treated as well as control animals.

### Assessment of Anti-proliferative effect of EpCAM-siRNA chimeras

To assess the anti-proliferative effect of the chimeras on the xenografts, immunohistological analysis for cell proliferation markers Ki67, p53 and anti-proliferative marker BCL2 was performed. H&E staining revealed lot of dead cells in EpCAM-siRNA treated conditions. Ki67, a nuclear specific marker for cell proliferation, was highly expressed in control mice compared to all the aptamer chimera treated tumour tissues (**Figure 6**). The decrease in Ki67 reactivity indicates decreased cancer cell proliferation after aptamer chimera treatments, suggesting the anti-proliferative role of the EpCAM-siRNA chimeras. We then tested for the anti-proliferative and anti-apoptotic markers, p53and BCL2, respectively, to understand if the tumour cell death is mediated through apoptosis as all the three silenced genes in the present study are found in p53 signalling pathway. After the administration of the aptamer chimeras, p53 protein expression increased and BCL2 protein expression decreased in EpCAM-siPLK1 & EpCAM-siSTAT3 treated mice compared to the control. For studying the proliferation after the EpCAM-BCL2 aptamer chimera treatment, the tumour tissue from the respondent mice was used for studying Ki67, P53 and BCL2 protein expression. In these tumours Ki67 and BCL2 protein expressions decreased with concomitant increase in P53. However non-responders had no upregulation of P53 (data not presented). The switching of proliferation of the tumour cells into apoptosis was brought after the aptamer chimera treatments. This ensures that the cells are undergoing apoptosis when a particular oncogene (PLK1, BCL2, STAT3) was silenced, resulting in the suppression of tumour growth.

**Figure 6:**
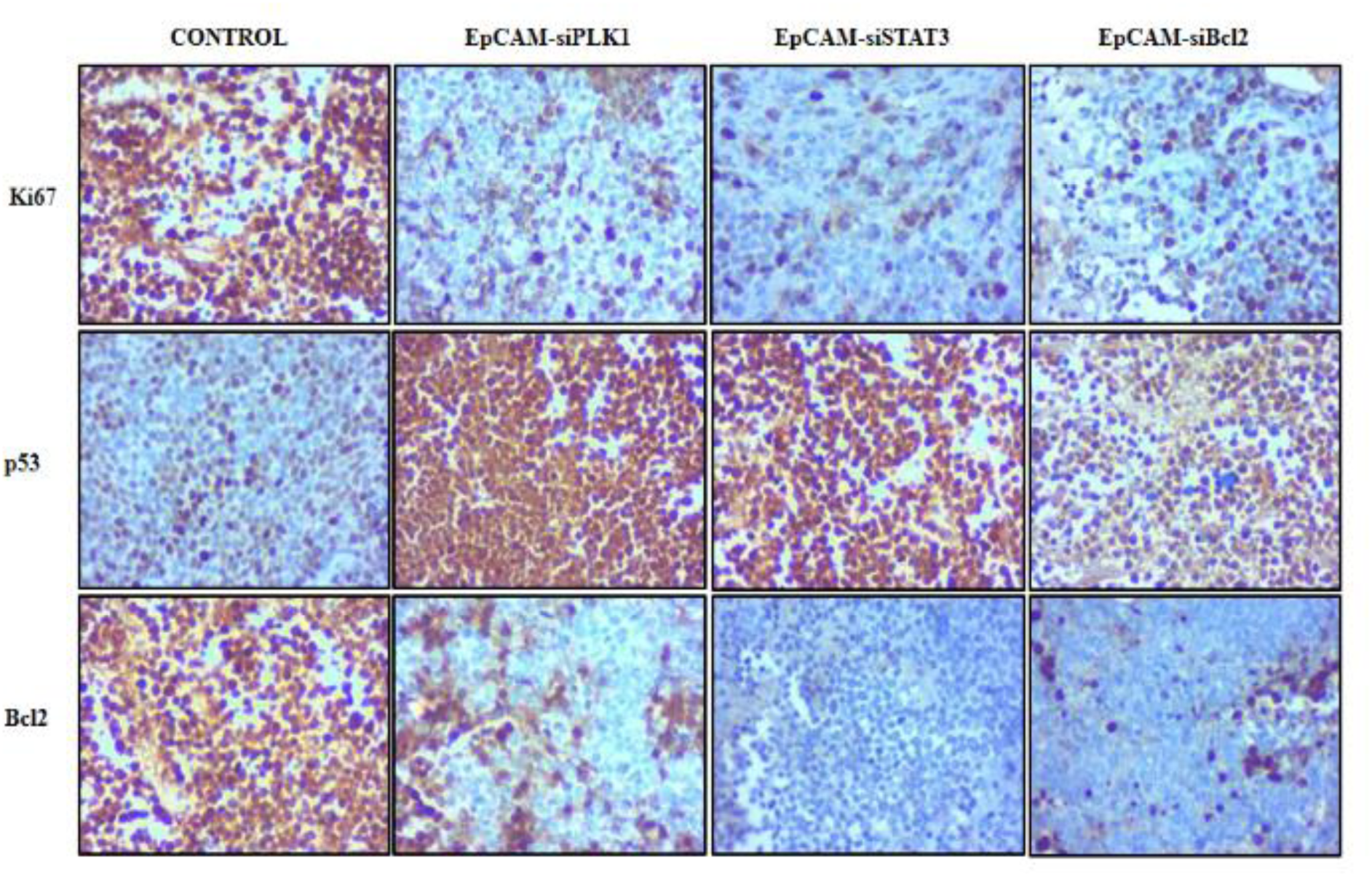
Evaluation of anti-proliferative and anti-apoptotic genes after EpCAM-siRNA treatment *in vivo*. formalin-fixed paraffin embedded sections of tumours were stained with antibodies targeting Ki67, p53 and BCL2. Compared to control, chimera treatment has significantly reduced the expression of Ki67 and BCL2, and up-regulated p53 (40x magnification).

### Phosphoproteomics identified altered signalling proteins

We observed that EpCAM-siPLK1 chimera exhibited higher therapeutic efficacy compared to the other two chimeras in both *in vitro* and *in vivo* models (**Figures 3b and 5**). PLK1, a kinase, plays a major role in phosphorylating many oncoproteins. Silencing PLK1 would certainly have an impact on the phosphorylation status of its downstream targets. Hence we chose to study the altered signalling pathways upon silencing PLK1 by Stable Isotope Labelling with Amino acids in Cell culture (SILAC)-based phosphoproteomics analysis. We identified about 3501 phosphopeptides PLK1-dependent phosphorylation sites which mapped to 1493 proteins (**Supplementary data 1**). We considered 1.5-fold change in phosphorylation status of identified peptides for analysing differential expression of proteins when PLK1 is down regulated. A set of 904 peptides were hypophosphorylated and 1382 were hyperphosphorylated after the EpCAM-siPLK1 aptamer chimera treatment, which corresponded to 552 hypophosphorylated and 798 hyperphosphorylated proteins. Partial list of hypophosphorylated and hyperphosphorylated proteins and the corresponding pathways are given in **Table 1**. Representative MS/MS spectra of peptides of the hyperphosphorylated and hypophosphorylated proteins in EpCAM-siPLK1 treated cells are shown in **FigureS3**. DAVID’s Functional annotation tool and KEGG pathway analysis identified cell cycle regulation, mTOR MAPK and Wnt signalling pathway as major deregulated pathways after the PLK1 knockdown. Additionally, the classic PLK1 cell cycle regulation through CDC25, p53 signalling pathway and AMPK regulation were the prominent pathways identified after the knockdown of PLK1 gene.

**Table 1:**
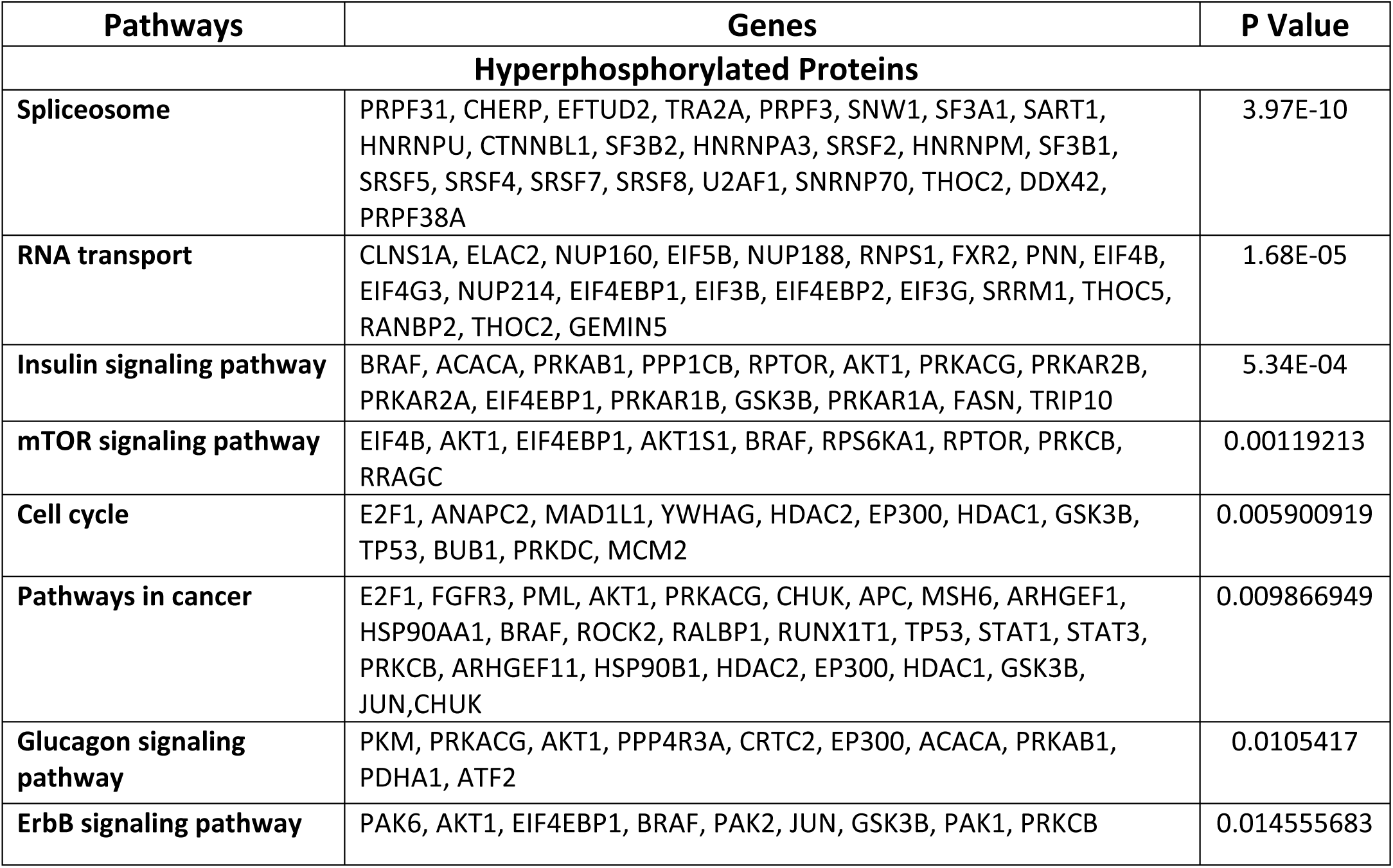
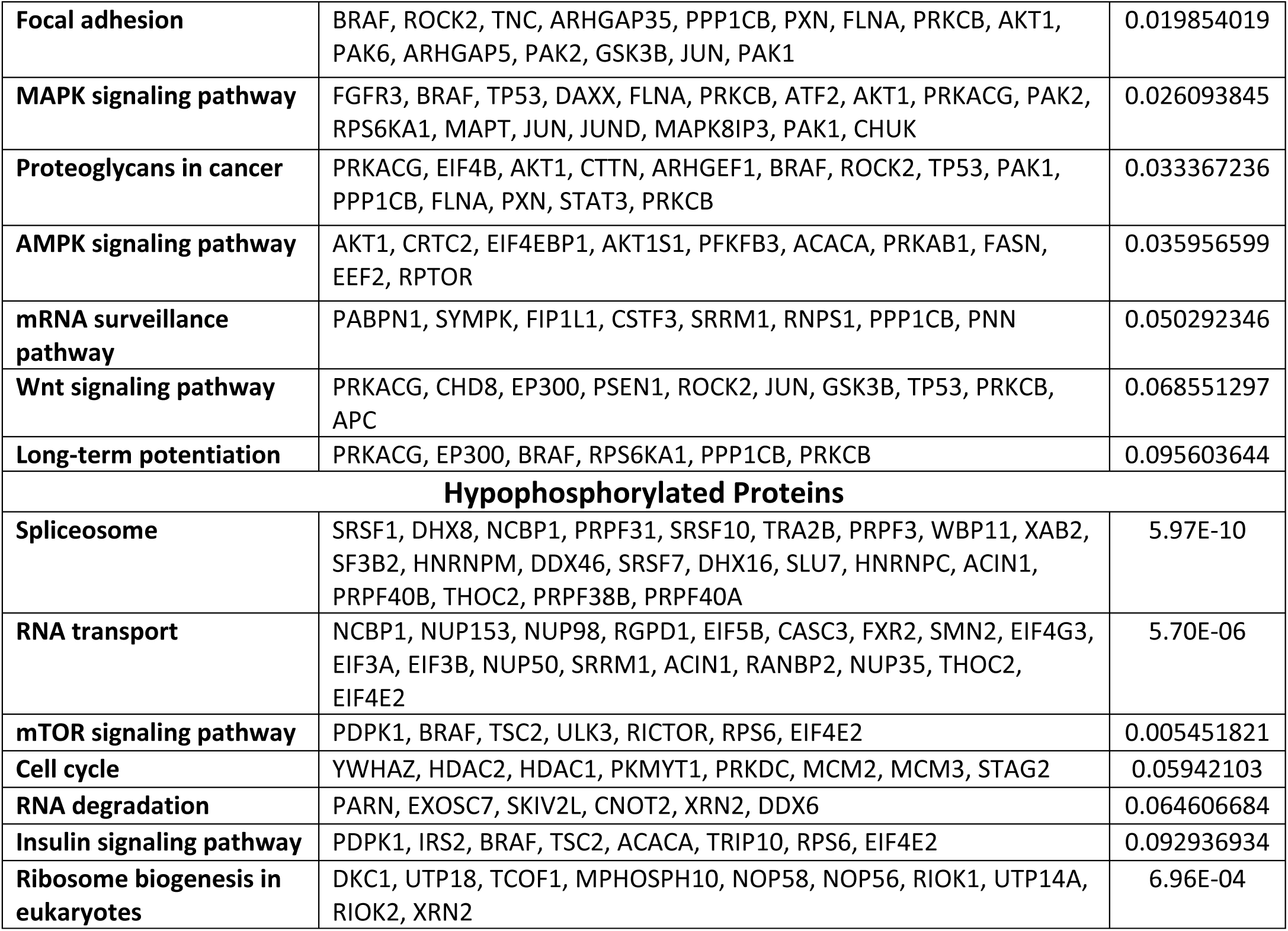
List of hyperphosphorylated and hypophosphorylated proteins and the respective pathways they are involved. The pathways were obtained by using DAVID’s functional annotation analysis.

Interestingly, p53 phosphorylation was identified in our global phosphoproteomic analysis (**Supplementary data 1**) and increased p53 protein expression was seen in EpCAM-siPLK1 treated mouse tumours (**Figure 6**). Therefore, we examined several phosphorylated proteins that are related to P53 signalling pathways. 14-3-3 proteins which regulate cell cycle were found to be downregulated **(Supplementary data 1)**. Additionally, the expressions of cell cycle regulators like Chk1, cdc25, p53 and cdk1 were altered after the PLK1 knockdown. Major proteins which are known to be responsible for p53 phosphorylation were also identified in the study. We found that DNA-PK, p300, CBP, GS3Kβ, IKK and APC/C as hyperphosphorylated whereas GTSE1 was hypophosphorylated after the silencing PLK1 stating the reason for enhanced levels of p53. We also found proteins involved in the cell survival pathway (PKB/Akt, GSK3β, IKK and APC/C) to be regulated after PLK1 knockdown. Interestingly, dysregulation of phosphorylation was also observed in the proteins associated with the metabolic regulation of the cells, where S6 and EIF4E-BP1 was hypophosphorylated and hyperphosphorylated, respectively. Additionally, AMPK was hyperphosphorylated when PLK1 is down regulated.

## DISCUSSION

Therapeutic application of aptamer-siRNA chimeras for treating cancer is a promising technique to deliver the siRNA to specific tumour cells and eliminate them completely while reducing the off target effects on the normal cells ^21^. Presently, we designed aptamer chimeras to specifically target cancer cells by recognising overexpression of EpCAM and deliver the siRNA that regulate the oncogene expression in these cells. In consistence with the designed features, the reduction in the oncogene expression in the cancer cells was dependent on the cellular EpCAM expression levels. The knockdown of the gene in the various cell lines was completely based on their EpCAM expression levels in a very cell-type specific manner. Higher EpCAM expression on MCF7, NCC RB 51 cells resulted in greater binding of the, EpCAM-siPLK1, EpCAM-siSTat3 and EpCAM-siBCL2 aptamer chimeras internalisation and effectively reducing the PLK1, STAT3 and BCL2 oncogene expressions, respectively, at lower aptamer treatments. A moderate EpCAM expression in HepG2 and RPMI 2650 needed higher dose of aptamer chimera concentration to elicit cellular toxicity. The control cell line, Mio M1, showed very minimal response to the three aptamer chimeras, whereas NCC RbC 51 showed the maximum ability to kill cancer cells. Additionally, *in-vivo* mouse xenograft model showed effective tumour reduction in nude mice on treatment with these chimeras. Several EpCAM conjugations like EpCAM-siRNA conjugations, EpCAM-drug and EpCAM-toxin conjugations have shown high efficacy in reducing cancer cell growth ^21, 22^. Present study indicates that EpCAM mediated silencing of PLK1, STAT3 and BCL2 oncogenes can show targeted therapeutic ability to reduce tumours in mouse models.

The therapy response in mouse model could be due to the increased expression of p53 tumour suppresser protein and decreased anti-apoptosis protein BCL2. EpCAM-siPLK1 exhibited superior functional ability by greater tumour burden reduction compared to other two aptamers. The signalling mechanism associated with superior therapeutic performance is further studied by global phosphoproteomics and mass-spectrometry. This technique is a valuable tool for identifying signalling after drug treatments in cancer models ^23, 24^. The Bioinformatic analysis of deregulated phosphoproteins after PLK1 down regulation identified altered Map kinase, cell cycle and Wnt signalling pathways. The role of PLK1 in cell cycle progression through map kinase pathway is further strengthened using our global phosphoproteomic analysis in RB cells. The cross talk and integration of these two pathways leading to cancer is proposed in several cancers^25^. Similarly an alteration of Wnt signalling after the down regulation of PLK1 could also account for observed therapeutic effect in RB cells. An increased PLK1 expression modulated Wnt signalling pathway was identified in prostate cancer^26^.

On silencing PLK1, the increased levels in p53 and phosphorylation might also be a major reason for cell death via apoptosis. Over expression of PLK1 is known to phosphorylate GTSE1 and Topors which are direct substrates of p53 leading to inactivation of p53 ^27, 28^. EpCAM-siPLK1 mediated p53 accumulation in our study could be due to the hypophosphorylation of GTSE1. Furthermore, p53 is also a downstream target of DNA-PK which is regulated by PLK1 during mitosis ^29^ and the presence of DNA-PK results in the inactivation of Akt/PKB and activation of GSK3β, ultimately contributing to the accumulation of p53^30^. Alternatively, the increased p53 expression mediated by GSK3β have also been reported to result in Cyclin D1 expression leading to apoptosis ^31^. Zhiguo Li showed that depletion of PLK1 inhibited the PI3K pathways ^32^ which coincide with our results where GS3Kβ, IKK, APC/C and p53 were hyperphorphorylated. Additionally, the increased phosphorylation seen in the tumour suppressor protein AMPK might be the strong reason for the metabolic shift in the functioning of the cells. The active form of PLK1 is known to be responsible for the phosphorylation of the downstream targets of AMPK such as S6 and 4E-BP1 through the mTOR pathway. Alternatively, p53 can also regulate the mTOR pathway through AMPK activation thereby increasing apoptosis in RB cells. Collectively, our phosphoproteomics analysis demonstrated that EpCAM-siRNA mediated knockdown of PLK1 eventually lead to alteration in cell cycle progression mediated through P53/ MAPK/Wnt signalling leading to apoptosis.

In conclusion, non-specificity of therapy in cancers is addressed using cell-type specific aptamer chimeras which kills the cancer cells alone. The EpCAM aptamer chimeras could effectively reduce tumours in mouse xenograft models. This work also sheds light on the importance of PLK1 in cancer, and silencing the gene could lead to apoptosis at various cell cycle check points. The reduction in cancer growth could be due to the upregulation of p53/MAPKinase/Wnt mediated cellular signalling leading to increased cancer cell death. Our study paves a platform for efficient second line of therapies in addition to existing chemotherapy options for controlling cancer cell proliferation, reducing the off-target effects on normal cells.

## MATERIALS AND METHODS

### Reagents

For cell culture, RPMI 1640, DMEM, antibiotic-antimycotic solution, and FBS were purchased from Gibco Life Technologies, Waltham, MA, USA.

### Aptamer design

Aptamer chimeras EpCAM-siPLK1, EpCAM-siSTAT3 and EpCAM-siBCL2 and EpCAM-siScr were designed. The lengths of the chimeras were designed to be 60 nucleotides containing a dicer recognition site between the EpCAM aptamer and the siRNA sequence. To increase the stability of the aptamer chimeras against endogenous nucleases, modifications like, adding 2 UU-nucleotide 3’ overhangs, optimizing the thermodynamic profile and structure of the duplex to favor the processing of siRNA guide strand, were made. The secondary structure prediction was done using RNA structure program version 5.0. The aptamer was fluorescently labelled with FITC at the 3’ end to visualize binding of chimeras to the cells. The designed aptamer-chimeras were custom synthesized through Dharmacon, Lafayette, CO, USA.

### Dicer cleavage assay

All EpCAM-siRNA chimera (100 nM) was denatured by heating for 10 min in a boiling water bath. Digestion using Dicer enzyme was set up as per the manufacturer’s protocol (Recombinant Dicer KitT520002, AMS Biotechnology, UK) with and without Dicer. The samples were then resolved using native PAGE (15% w/v polyacrylamide) and stained with ethidium bromide before visualization using the Gel Doc XR System (Bio-Rad).

### Cell culture

NCC Rb C 51RB cells (obtained from RIKEN BioResource Center, Ibaraki, Japan) were maintained in RPMI 1640 (Roswell Park Memorial Institute 1640), with 10% Fetal bovine serum (FBS) and supplemented with antibiotic-antimycotic solution containing penicillin 100 µg/ml, streptomycin 100 µg/ml, and amphotericin B 250 ng/ml. Breast cancer cell line MCF-7 (Michigan Cancer Foundation-7) was procured from the American Type Culture Collection (ATCC), Manassas, VA, USA, and Müller Glial Mio M1 cell line was a kind gift from Professor Astrid Limb, London, UK, and they were cultured in DMEM (Dulbecco’s Modified Eagle’s Medium) containing 10% FBS with and without antibiotic-antimycotic solution, respectively. RPMI 2650 (Head and Neck cancer cell line) and HepG2 (Hepatocellular cell line) were procured from NCCS, Pune, India and were cultured in αMEM (Dulbecco’s Modified Eagle’s Medium) containing 10% FBS with antibiotics. All the cell lines were maintained at 5% CO_2_ environment at 37 °C.

### Transfection

At about 85% confluency, the cells were treated with the above mentioned siRNA and EpCAM-siRNA chimeras. 100 nM of siRNA along with lipofectaminefor 24 h was used for transfection for all the cell lines. In the case of EpCAM-siRNA chimeras different concentrations ranging from 100-500 nM was used depending on the cell line for 48 h.

### Flow cytometry for binding efficacy

Approximately, 4×10^5^cells (Mio M1, NCC RbC 51, MCF-7, RPMI 2650, HepG2) were collected and washed twice with cold phosphate-buffered saline (PBS). Cells were then resuspended in blocking buffer (PBS with 0.02% sodium azide, 0.1% FBS, and 0.01% Triton X-100) and incubated with different concentrations (100 nM - 500 nM) of FITC tagged aptamer chimeras (EpCAM-siPLK1, EpCAM-siSTAT3 and EpCAM-siBCL2). After 1 h, the cells were washed twice with ice cold PBS and resuspended in sheath fluid. About 1×10^4^ cells were collected and analysed for binding efficacy of the chimeras using FACS (BD FACS Calibur system).

### Flow Cytometry to check apoptosis using ANNEXIN/PI kit

Annexin/Propidium Iodide staining kit from BD Pharmingen, India (cat no:556547) was used to check the apoptotic levels. About 4×10^5^cells (Mio M1, NCC RbC 51, MCF-7, RPMI 2650, HepG2) were collected and washed twice with cold phosphate-buffered saline (PBS). Cells were then resuspended in 100 µl of 1X binding buffer present in the kit and incubated with 3 µl of Annexin and 2.5 ul of Propidium iodide. Propidium iodide alone and Annexin alone stained samples were used for comparison. After 1 h, the cells were washed twice with 1X Binding buffer, resuspended in sheath fluid, and analysed for apoptotic effect of the chimeras using FACS (BD FACS Calibur system).

### MTT assay

The cells (Mio M1, NCC RbC 51, MCF-7, RPMI 2650, HepG2) were treated with 100 nM - 500 nM of the chimeras for 72 h. Cell proliferation was checked by incubation with MTT (5 mg/ml) for 4 h at 37 °C. After 4 h, the formazan crystals were formed and upon addition of DMSO, the purple colour of formazan was observed. The absorbance was measured at 570 nm on ELISA plate reader (SpectraMaxPlus M4 Microplate reader).

### Quantitative RT-PCR

Mio M1, RPMI 2650 and HepG2 cells were transfected with 300 nM EpCAM-siRNA and NCC RbC 51 and MCF-7 cells were transfected with 100 nM EpCAM-siRNA. Cells treated with 100 nM of siPLK1, siSTAT3 and siBCL2using Lipofectamine 2000 reagent served as transfection controls and untreated cells were considered as controls. The cells were harvested after 48 hrs and washed twice with cold PBS. RNA was extracted with TRIzol reagent using an optimized protocol and reverse transcribed into cDNA using High-Capacity cDNA Reverse Transcription Kit (Applied Biosystems). Quantitative RT-PCR was performed with SYBR Green (Applied Biosystems ^®^ 7500) using GAPDH as endogenous control. The primers used were FP (5’-GACAAGTACGGCCTTGGGTA), RP (5’-GTGCCGTCACGCTCTATGTA) for PLK1 (Sigma-Aldrich), FP (5’-ATGTGTGTGGAGAGCGTCAA), RP(5’ACAGTTCCACAAAGGCATCC) for BCL2 (Sigma-Aldrich), FP (5’ACCTGCAGCAATACCATTGAC), RP (5’AAGGTGAGGGACTCAAACTGC) for STAT3 (Sigma-Aldrich) and FP (5’-AGAAGGCTGGGGCTCATTTG), RP (5’-AGGGGCCATCCACAGTCTTC) for GAPDH (Sigma-Aldrich). The threshold cycle (Ct) value obtained from Applied Biosystems were used to calculate relative quantification with the formula 2^−ΔΔCt^, where ΔΔCt are the values obtained on normalizing the values of treated samples with values of untreated and endogenous controls.

### Western blotting

Mio M1, RPMI 2650 and HepG2 cells were transfected with 300 nM EpCAM-siRNA and NCC RbC 51 and MCF-7 cells were transfected with 100 nM EpCAM-siRNA. The cells were then harvested after 48 h and lysed in RIPA buffer containing Protease Inhibitor Cocktail (Sigma-Aldrich) followed by cold PBS wash. Supernatant was collected after centrifuging the samples for 10 min at 10000 rpm. The protein concentration was estimated using Pierce™ BCA Protein assay kit (Cat no:23225), Thermofisher scientific, Waltham, MA, USA) method against BSA as a standard. 75 µg protein was resolved on a 10% SDS PAGE and transferred onto a nitrocellulose membrane. The blot was blocked for 1 h with 5% skimmed milk and the membrane was probed with anti-PLK1 primary monoclonal antibody (Invitrogen: 37-7000), anti-BCL2 primary monoclonal antibody (Invitrogen: 138800) and anti-STAT3 primary monoclonal antibody (Cell Signalling: 124H6) at 1:1000 dilution. The primary antibody was treated overnight at 4 °C. Then the blot was washed 3 times with PBST and probed with secondary anti-mouse conjugated with HRP (Santa Cruz Biotechnology: sc-2005) at 1:4500 dilution for 1 h. The membrane was detected using a TMB substrate (Merck). GAPDH (Sigma-Aldrich: G8795) was used as endogenous control.

### Statistical analysis

For quantitative RT-PCR and western blotting experiments, data obtained from triplicates were analysed using Statistical Package for the Social Sciences (SPSS) software. The comparisons were performed using Student’s *t*-test. *P* values less than 0.05 were considered as statistically significant.

### NCC RbC 51 xenograft study

#### Culture of Tumour Cells

Xenograft studies were carried out at Biology Division, Syngene International, Bangalore, India. All procedures were performed in a laminar air flow hood following sterile techniques. Human NCC RbC 51, RB cells with a viability of >90 % was chosen for the study. Approximately 5 × 10^6^ NCC RbC 51cells were re-suspended in 200 µL of serum-free media containing 50% of matrigel and kept on ice until cell injection.

#### Subcutaneous Injection of Cells

Male athymic nude mice (Hsd: Athymic Nude-Foxn1^nu^) housed in Individually Ventilated Cages (IVCs) were used for the present investigation. Cells were propagated in animals by injecting the cells subcutaneously in the right flank of the animals. The implanted area was monitored for tumour growth. Once the tumour attained a palpable size, animals were randomized based on tumour volume (TV ≈ 200 mm^3^) and dosing was initiated.

#### Formulation and Drug Dosage

Preparation of the test compound and dosing solutions

Required quantity of aptamers (Anti-PLK1, Anti-BCL2 and Anti-STAT3) were diluted in sterile water for injection to get final concentration of 8 nmol/ml as a working solution to deliver dose of 0.8 nmol/100µl/animal. Aliquots were prepared as per daily required doses and stored at −20°C. One tube from each test compound was thawed daily just prior to dose administration. The dose volume was maintained constant at 100 µL/animal. A single dose regimen was planned (Schedule −1: Q2D X 2 weeks-day 0, 2, 4, 6, 8, 10, 12 & 14).

#### Xenograft mouse model

EpCAM-siPLK1, EpCAM-siSTAT3, EpCAM-siBCL2 was intraperitoneally injected into the mice every third day for 30 days and a total ten cycles were carried out. Tumour sizes and body weights were measured twice a week. The tumour volume was calculated according to the formula: TV(mm^3^) = (L × W^2^)/2 (W, the width; L, length). The animals were euthanized two days after the last treatment. The tumours and organs were collected and fixed in 10% formalin buffer. The sections of tissues were analysed by H&E staining and immunohistochemistry.

#### Histology assay

Animals were euthanized with CO_2_, and tumours were collected for all the four conditions. The vital organs (Liver, Kidney, lymph nodes, heart, lung, spleen and brain) were removed and fixed with 4% paraformaldehyde in 0.1 M sodium phosphate buffer (pH 7.6). The serum was also collected separately. Tissues were cut into 10 mm sections and were dehydrated in alcohol-xylene, and embedded in paraffin. Sections were cut and mounted on to glass slides, deparaffinised in xylene and ethyl alcohol. Each block has a section for H&E staining. For immunohistochemistry assay, sections were incubated in 3% normal goat serum for 2 h and followed by overnight incubation with primary antibodies: Ki67, P53, BCL2, Cyclin D1(1:1000). After washing, the sections were incubated with biotinylated secondary antibody (1:200, VECTOR, Burlingame, CA) for 1 h. Following washing, the sections were incubated with VECTASTAIN ABC reagents for 30 min. The immunoreactivity (IR) was visualized with the substrate solution (VECTOR). The images were captured with Nuance fluorescence microscope with bright field imaging system.

#### SILAC Labelling, trypsin digestion and TiO 2 based phosphoprotein enrichment

2*10 6 cells were grown and incorporated with radio labelled isotopes. RPMI with 10% fetal bovine serum (FBS) supplemented with Antibiotic was used as the medium for both Heavy and light labelled conditions. Those incorporated with 13 C 6 -Lysine, 13 C 6 -Arginine were termed as heavy labelled cells, and those with normal lysine arginine were termed as light labelled cells. The cells were grown in the heavy/light media for about 6 passages, till the labelling was incorporated. The Light labelled cells were treated with EpCAM-siPLK1 and incubated for 48 h. After treatment the cells were lysed with 2% SDS. Equal amount of Heavy and Light labelled lysates were mixed together and mixed protein lysate was reduced with 5mM DTT at 60 o C for 30 minutes and alkylated using IAA at room temperature for 10 minutes. Protein precipitation was done using 6 volume of ice cold acetone and digested overnight at 37 o C using modified trypsin (Promega). Then peptides were fractionated by bRPLC and 8 fractions were collected. TiO2-based phosphopeptide enrichment method was used to enrich the phosphopeptides from each fraction. Then enriched peptides were dried and cleaned using C18 stage tips.

#### LC-MS/MS analysis and data analysis

The peptides were analysed on a Thermo Scientific™Orbitrap Fusion™ Tribrid™ Mass Spectrometer (Thermo Scientific, Bremen, Germany) interfaced with Easy-nLC 1000II nanoflow liquid chromatography system (Thermo Scientific, Odense, Denmark). Dried peptides were reconstituted in 0.1% formic acid and loaded on trap column (5µm × 2cm, packed in-house with Magic C18 AQ (MichromBioresources, Inc., Auburn, CA, USA). The peptides were then resolved on analytical column (75µm × 25cm, packed with 3µm ReproSilpur C18 AQ) at a flow rate of 280 nL/min using a gradient from 3 to 30% of solvent B (95% ACN and 0.1% formic acid) over 105 minutes. MS data were acquired using scan range of 400-1600 m/z at mass resolution of 120K and MS/MS data were acquired using resolution of 30,000. The instrument was operated for cycle time of 3 sec. Top abundant ions were isolated and fragmented using HCD with isolation width of 2 m/z and normalized collision energy of 34%. Mass spectrometry data was searched through the Proteome Discoverer platform (v2.1, Thermo Scientific, Bremen, Germany) using Mascot and SEQUEST search algorithms against Human RefSeq75 protein database supplemented with frequently observed contaminants. The search parameters for both the algorithms included maximum of two missed cleavage, carbamidomethylation at cysteine and SILAC labels 13C6-Lysine; 13C6-Arginine as fixed modification. Oxidation at methionine and phosphorylation at serine, threonine and tyrosine were set as dynamic modification. The data was searched against decoy database and false discovery rate was set <1% at PSM level. The probability of phosphorylation at S/T/Y site on each peptide was calculated using PhosphoRS node and peptides with more than 75% phosphoRS probability were considered to fetch the phospho site on protein level. The KEGG pathways were obtained using DAVID’s functional annotation bioinformatics analysis. All MS data have been deposited in the PRIDE database with identifier PXD008221.

## Supporting information

Supplementary Table 1: Phosphopeptides regulated by PLK1

## ACKNOWLEDGEMENTS

This work was funded by Department of Biotechnology (DBT), Govt. of India project entitled “To engineer aptamer siRNA chimeric conjugates for targeting epithelial cancers” (BT/PR3580/PID/6/625/2011) and DBT-COE project (BT/01/CEIB/11/V/16).

## AUTHOR CONTRBUTIONS

JB participated in designing and conducting the experiments and interpretation of the data. LB provided assistance in conducting the experiments. BC conducted the proteomic experiments. HG and AP supervised the proteomic analysis. KS and LBS participated in the study design and giving important feedback. SVE conceived the project. JB and SVE wrote the paper. All the authors edited and approved the project.

## REFERENCES

1. Mustonen, E.K., Palomäki, T. & Pasanen, M. (2017). Oligonucleotide-based pharmaceuticals: Nonclinical and clinical safety signals and non-clinical testing strategies. Regulatory Toxicology and Pharmacology. 90: 328–341

2. Kruspe, S. & Giangrande, P.H. (2017). Design and Preparation of Aptamer-siRNA Chimeras (AsiCs) for Targeted Cancer Therapy’, Methods in Molecular Biology. 1632: 175–186

3. Lakhin A.V., Tarantul V.Z., Gening L.V. Aptamers: Problems, solutions and prospects. ActaNaturae. 2013;5:34–43

4. Brody E.N., Gold L. Aptamers as therapeutic and diagnostic agents. J. Biotechnol. 2000;74:5–13. doi:10.1016/S1389-0352(99)00004-5

5. Zhou, J., Bobbin, M.L., Burnett, J.C. & Rossi, J.J. (2012). Current progress of RNA aptamer-based therapeutics. Frontiers in Genetics. 3:234

6. Shigdar, S., Lin, J., Yu, Y., Pastuovic, M., Wei, M. & Duan, W. (2011). RNA aptamer against a cancer stem cell marker epithelial cell adhesion molecule. Cancer Science. 102(5):991–8

7. McNamara, J.O., Andrechek, E.R., Wang, Y., Viles, K.D., Rempel, R.E., Gilboa, E., et al. (2006). Cell type-specific delivery of siRNAs with aptamer-siRNA chimeras. Nature Biotechnology. 24(8):1005–15

8. Zhou, J., Tiemann, K., Chomchan, P., Alluin, J., Swiderski, P., Burnett, J., et al. (2013). Dual functional BAFF receptor aptamers inhibit ligand-induced proliferation and deliver siRNAs to NHL cells. Nucleic Acids Research. 41(7):4266–4283

9. Ranji, P., Salmani, K.T., Saeedikhoo, S. & Alizadeh, A.M. (2016). Targeting cancer stem cellspecific markers and/or associated signaling pathways for overcoming cancer drug resistance. Tumour Biology: The Journal of International society for Onco developmental Biology and Medicine. 37(10): 13059–13075

10. Mohammad, Z.K., Prachi, P., Rajiv, P.G. (2013). Role of STAT3 in Cancer Metastasis and Translational Advances. Biomed Res Int. 2013; 2013: 421821

11. Park, J.E., Hymel, D., Burke, T.R. & Lee, K.S. (2017). Current progress and future perspectives in the development of anti-polo-like kinase 1 therapeutic agents. F1000Research. 6:1024.

12. Xiong, A., Yang, Z., Shen, Y., Zhou, J. & Shen, Q. (2014). Transcription Factor STAT3 as a Novel Molecular Target for Cancer Prevention. Cancers (Basel). 6(2):926–57.

13. Nigg, E.A., Blangy, A. & Lane, H.A. (1996). Dynamic changes in nuclear architecture during mitosis: on the role of protein phosphorylation in spindle assembly and chromosome segregation. Experimental Cell Research. 229:174–180.

14. Kumagai, A. & Dunphy, W.G. (1996). Purification and molecular cloning of Plx1, a Cdc25regulatory kinase from Xenopus egg extracts. Science. 273(5280):1377–80.

15. Borner, C. (2003). The Bcl-2 protein family: sensors and checkpoints for life-or-death decisions. MolImmunol. 39(11):615–47.

16. Kirkin, V.,Joos, S. & Zörnig, M. (2004). The role of Bcl-2 family members in tumorigenesis. BiochimBiophysActa. 1644(2-3):229–49.

17. Singh, L., Pushker, N., Saini, N., Sen, S., Sharma, A., Bakhshi, S., Chawla, B. & Kashyap, S. (2015) Expression of pro-apoptotic Bax and anti-apoptotic Bcl-2 proteins in human retinoblastoma. ClinExpOphthalmol. 43(3):259–67.

18. Gil-Gómez, G., Berns, A., Brady, H.J. (1998). A link between cell cycle and cell death: Bax and Bcl-2 modulate Cdk2 activation during thymocyte apoptosis. The EMBO Journal. 17(24):7209–7218.

19. Siveen, K.S., Sikka, S., Surana, R., Dai, X., Zhang, J., Kumar, A.P, et al. (2014). Targeting the STAT3 signaling pathway in cancer: Role of synthetic and natural inhibitors. BiochimicaetBiophysicaacta. 1845(2):136–54.

20. Gilboa-Geffen, A., Hamar, P., Le, M.T., Wheeler, L.A., Trifonova, R., Petrocca, F. (2015). Gene Knockdown by EpCAM Aptamer-siRNA Chimeras Suppresses Epithelial Breast Cancers and Their Tumor-Initiating Cells. Molecular Cancer Therapeutics. 14(10):2279–91.

21. Subramanian, N., Raghunathan, V., Kanwar, J.R., Kanwar, R.K., Elchuri, S.V., Khetan, V. (2012) Target-specific delivery of doxorubicin to RB using epithelial cell adhesion molecule aptamer. Molecular vision. 18:2783–95

22. Jayashree, B., Srimany, A., Jayaraman, S., Bhutra, A., Janakiraman, N., Chitipothu, S. (2016). Monitoring of changes in lipid profiles during PLK1 knockdown in cancer cells using DESI MS. Anal Bioanal Chem. 408:5623–5632.

23. Van der Mijn JC, Broxterman HJ, Knol JC, Piersma SR, De Hass RR, dekker H et al (2016). Sunitinib activates Axl signaling in renal cell cancer. Int J cancer. 138 (12):3002–10.

24. Bai Y, Kim JY, Walter JM, Fanqb, Kinose F, Song L et al (2014). Adaptive responses to dasatinib-treated lung squamous cell cancer cells harboring DDR2 mutations. Cancer Res 74(24):7217–7228

25. Malumbres M, Barbacid. To cycle or not to cycle: a critical decision in cancer (2001). Nat Rev Cancer. 1(3):222–31

26. Cristobal L, Rojo F, Madoz-Gurpide, Gracia-Foncillas (2016). Cross Talk between Wnt/β-Catenin and CIP2A/Plk1 Signaling in Prostate Cancer: Promising Therapeutic Implications. Mol Cell Biol. 31:36(12):1734–9.

27. Subramanian, N., Kanwar, J.R., Kanwar, R.K., Sreemanthula, J., Biswas, J., Khetan, V. (2015). EpCAM Aptamer-siRNA Chimera Targets and Regress Epithelial Cancer. PlOS One. 10(7).

28. Liu, X.S., Li, H., Song, B., Liu, X. (2010). Polo-like kinase 1 phosphorylation of G2 and S-phase-expressed 1 protein is essential for p53 inactivation during G2 checkpoint recovery. EMBO Rep. 11(8):626–32.

29. Yang, X., Li, H., Deng, A., Liu, X. (2010). phosphorylation of Topors is involved in its degradation. MolBiol Rep. 37(6): 3023–3028.

30. Douglas, P., Ye, R., Trinkle-Mulcahy, L., Neal, J.A., De Wever, V., Morrice, N.A. (2014). Polo-like kinase 1 (PLK1) and protein phosphatase 6 (PP6) regulate DNA-dependent protein kinase catalytic subunit (DNA-PKcs) phosphorylation in mitosis. Biosci Rep. 34(3).

31. Boehme, K.A., Kulikov, R., Blattner, C. (2008). p53 stabilization in response to DNA damage requires Akt/PKB and DNA-PK. Proc Natl AcadScu U S A. 105(22):7785–90.

32. Li, Z., Li, J., Bi, P., Lu, Y., Burcham, G., Elzey, B.D. (2014). Plk1 Phosphorylation of PTEN Causes a Tumor-Promoting Metabolic State. Mol Cell Biol. 34(19):3642–61

