## Supplementary Figures for "EpCAM Aptamer siRNA chimeras: Therapeutic efficacy in epithelial cancer cells"

**SUPPLEMENTARY INFORMATION**

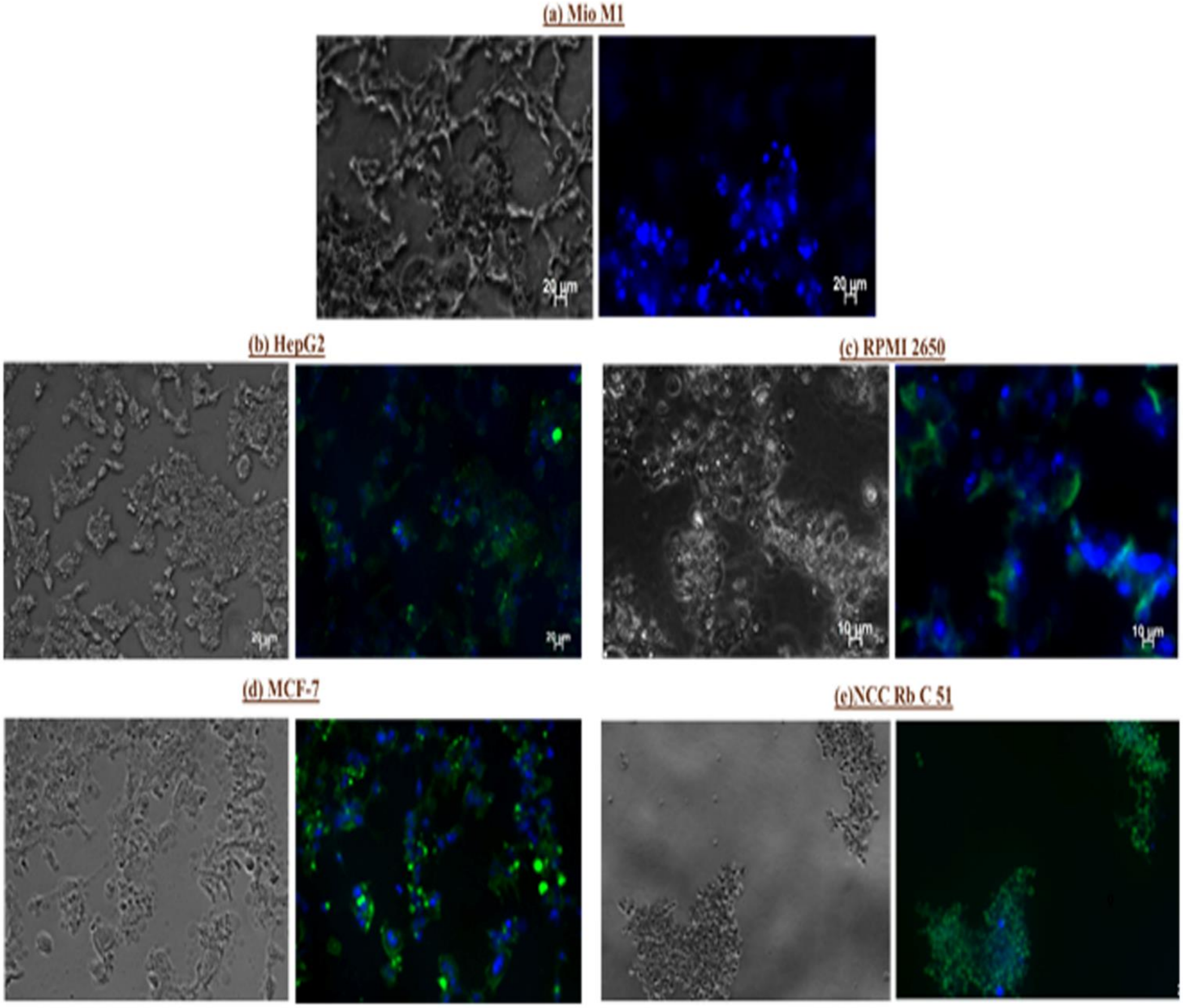


**S1: Aptamer-siRNA chimeras bind specifically to the cell surface antigen EpCAM. Cell lines** (a) Mio M1 (b) HepG2 (c) RPMI 2650 (d) MCF-7 and (e) NCCRbC 51were incubated with FITC labelled EpCAM-siRNA chimera and analysed by microscopy

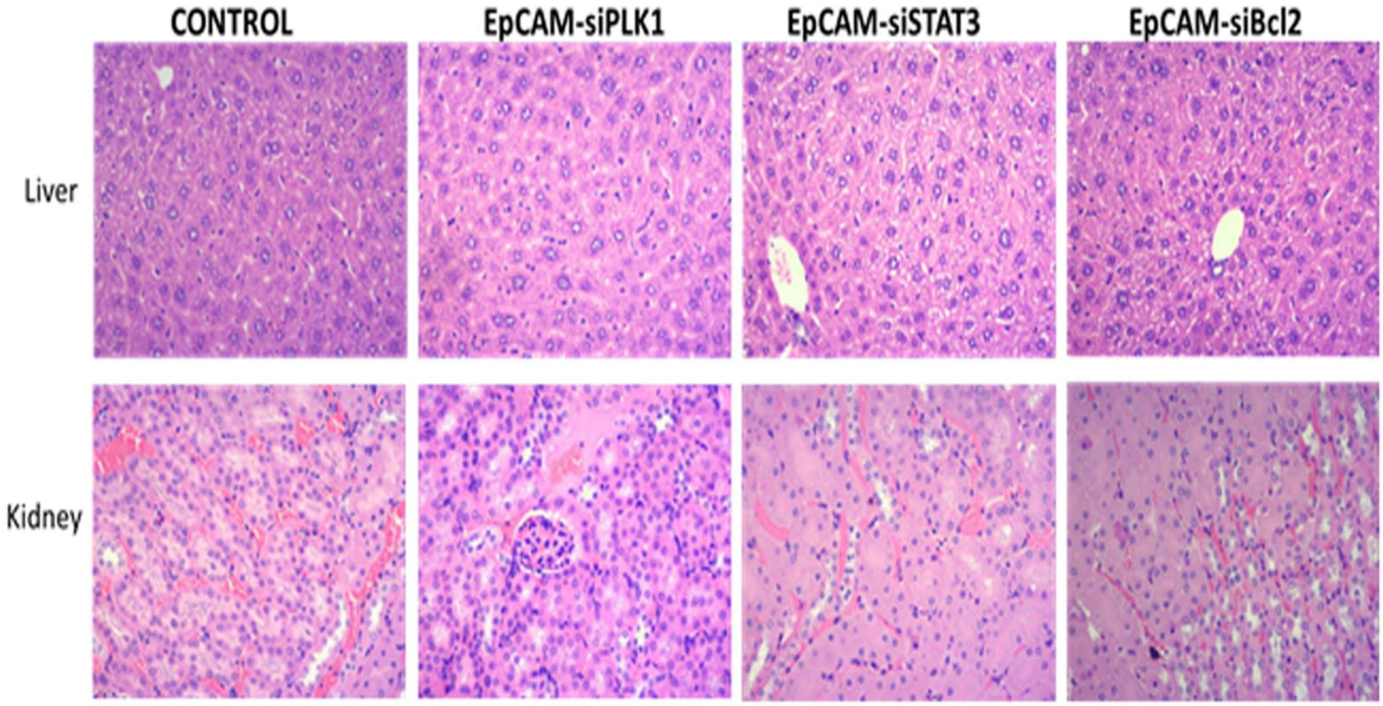


**Figure S2: H&E staining of the vital organs Liver and Kidney.** The morphology of these tissues remained unaffected after the chimera treatment.

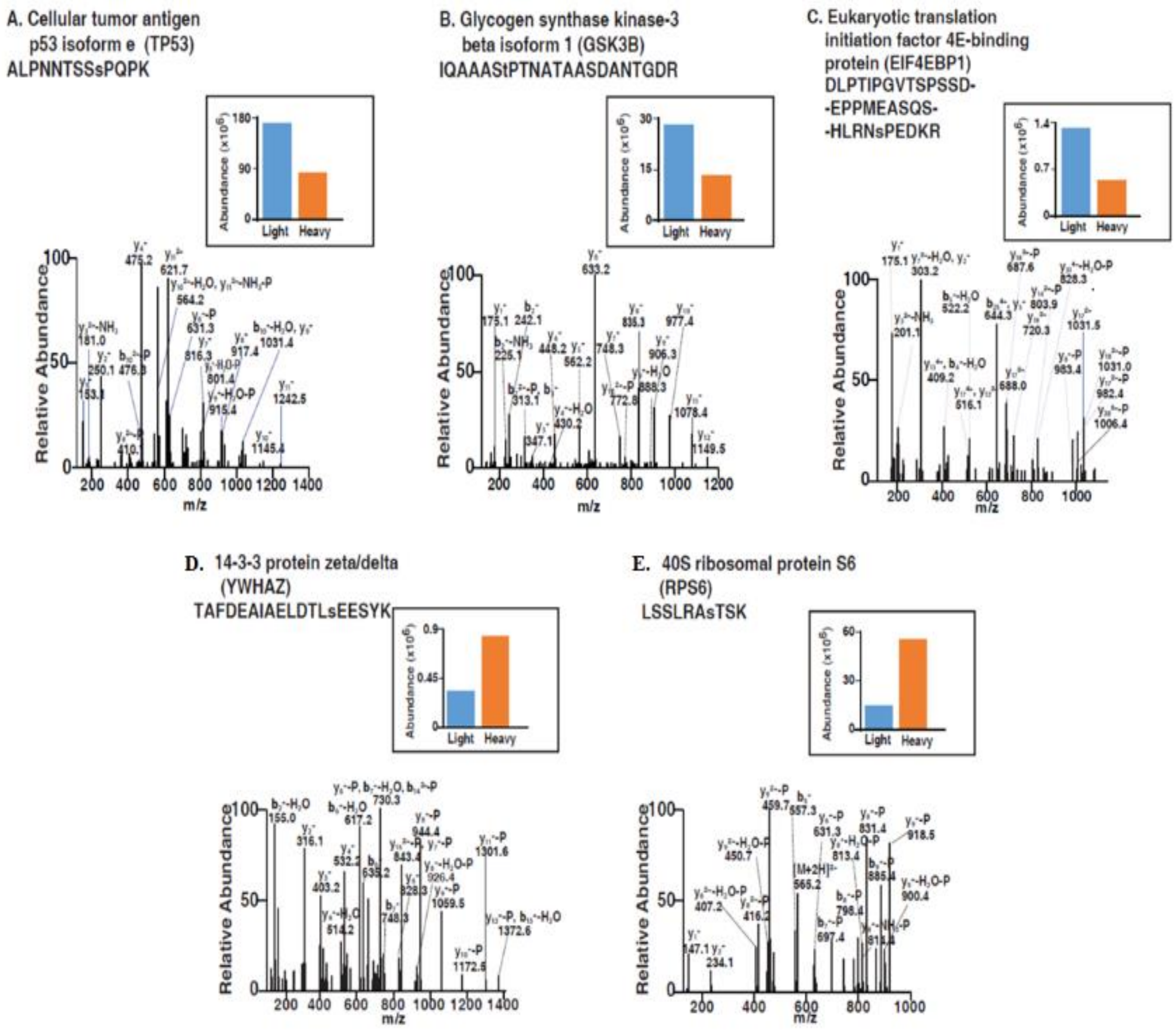


**Figure S3: Representative MS/MS spectra of peptides of hyperphosphorylaed proteins in EpCAM-siPLK1 treated cells:** (A) Cellular tumour antigen p53 isoform e (TP53), (B) Glycogen synthase kinase-3 beta isoform 1 (GSK3B) and (C) Eukaryotic translation initiation factor 4E-binding protein (EIF4EBP1). Representative MS/MS spectra of peptides of hypophosphorylaed proteins in EpCAM-siPLK1 treated cells: (D) 14-3-3 protein zeta/delta (YWHAZ) and (E) 40S ribosomal protein S6 (RPS6)
